# Inter-individual genomic heterogeneity within European population isolates

**DOI:** 10.1101/581470

**Authors:** Paolo Anagnostou, Valentina Dominici, Cinzia Battaggia, Stefania Sarno, Alessio Boattini, Carla Calò, Paolo Francalacci, Giuseppe Vona, Sergio Tofanelli, Miguel G. Vilar, Vincenza Colonna, Luca Pagani, Giovanni Destro Bisol

**Author notes:** These authors contributed equally to this work (Paolo Anagnostou, Valentina Dominici).

## Abstract

A number of studies carried out since the early ‘70s has investigated the effects of isolation on genetic variation within and among human populations in diverse geographical contexts. However, no extensive analysis has been carried out on the heterogeneity among genomes within isolated populations. This issue is worth exploring since events of recent admixture and/or subdivision could potentially disrupt the genetic homogeneity which is to be expected when isolation is prolonged and constant over time. Here, we analyze literature data relative to 87,818 autosomal single-nucleotide polymorphisms, which were obtained from a total of 28 European populations. Our results challenge the traditional paradigm of population isolates as genetically (and genomically) uniform entities. In fact, focusing on the distribution of variance of intra-population diversity measures across individuals, we show that the inter-individual heterogeneity of isolated populations is at least comparable to the open ones. More in particular, three small and highly inbred isolates (Sappada, Sauris and Timau in Northeastern Italy) were found to be characterized by levels of this parameter largely exceeding that of all other populations, possibly due to relatively recent events of genetic introgression. Finally, we propose a way to monitor the effects of inter-individual heterogeneity in disease-gene association studies.

## Introduction

Studying groups subject to barriers to gene flow provides a unique opportunity to understand how inbreeding and drift have shaped the structure of human genetic diversity. A very large number of investigations carried out since early ‘70s has examined the effects of isolation on intra- and inter-population variation in diverse geographical contexts, using genetic polymorphisms varying in mode of inheritance and evolutionary rate [e.g. 1–5]. Currently, the consequences of isolation may be better studied using genome wide approaches (GWAs), such as those based on single-nucleotide polymorphism (SNP) microarrays, which enable the simultaneous analysis of markers distributed across the human chromosomes. Compared to unilinearly transmitted polymorphisms of mitochondrial DNA and Y chromosome or to small panels of autosomal loci, GWA approaches make it possible to detect the imprints of isolation left on genomic makeup not only by mutation, but also by recombination [6–14].

In a previous study, we have compared intra and inter-population measures of genomic variation in a large sampling of European populations in order to understand to what extent the discrete open and isolated dichotomous categories correspond to the way in which their genomic diversity is structured [15]. Here, we move our focus to the heterogeneity among genomes within populations. Our results shed light on not yet understood aspects of the genomic structure of population isolates, which may also have significant implications for their use in disease-gene association studies.

In this study, we focus on the variance of intra-population variation measures in a large sampling of European populations using 87,818 autosomal SNP data. Our results highlight the existence of different and partly unexpected patterns, whose implications for our current view of the genetic structure of population isolates and disease-gene association studies are discussed.

## Materials and methods

### Dataset

We assembled a total of 87,818 autosomal SNPs, included in the GenoChip 2.0 array [16], in 610 healthy unrelated adult individuals from 28 European populations (Table 1). Our dataset comprises nine populations with clear signatures of genetic isolation evaluated using both unilinear and autosomal polymorphisms [15,17,18] plus nineteen open populations which were chosen using the following three criteria: (i) geographic proximity with the isolated populations; (ii) geographic coverage of the European continent; (iii) sample size of at least 10 individuals. Compared to the dataset used by Anagnostou et al. [15], we included five open populations (Belarus, Hungary, Lithuania, Romania and Ukraine) and removed the Cimbrians since it lacked consistent signatures of genetic isolation. Despite its limits [15], we maintain here the dichotomy between open and isolated population for practical reasons (see also the Discussion section).

**Table 1.**
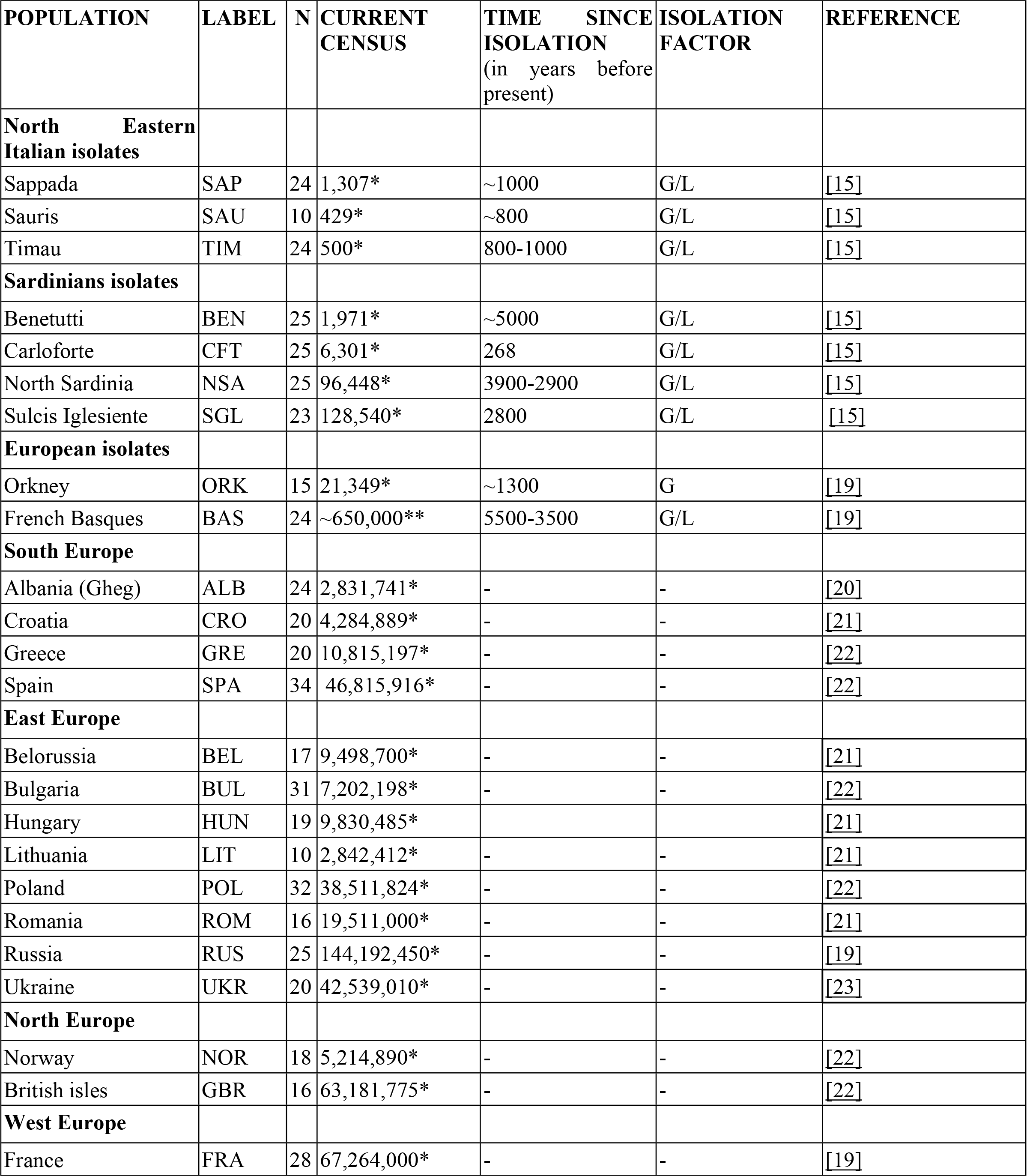
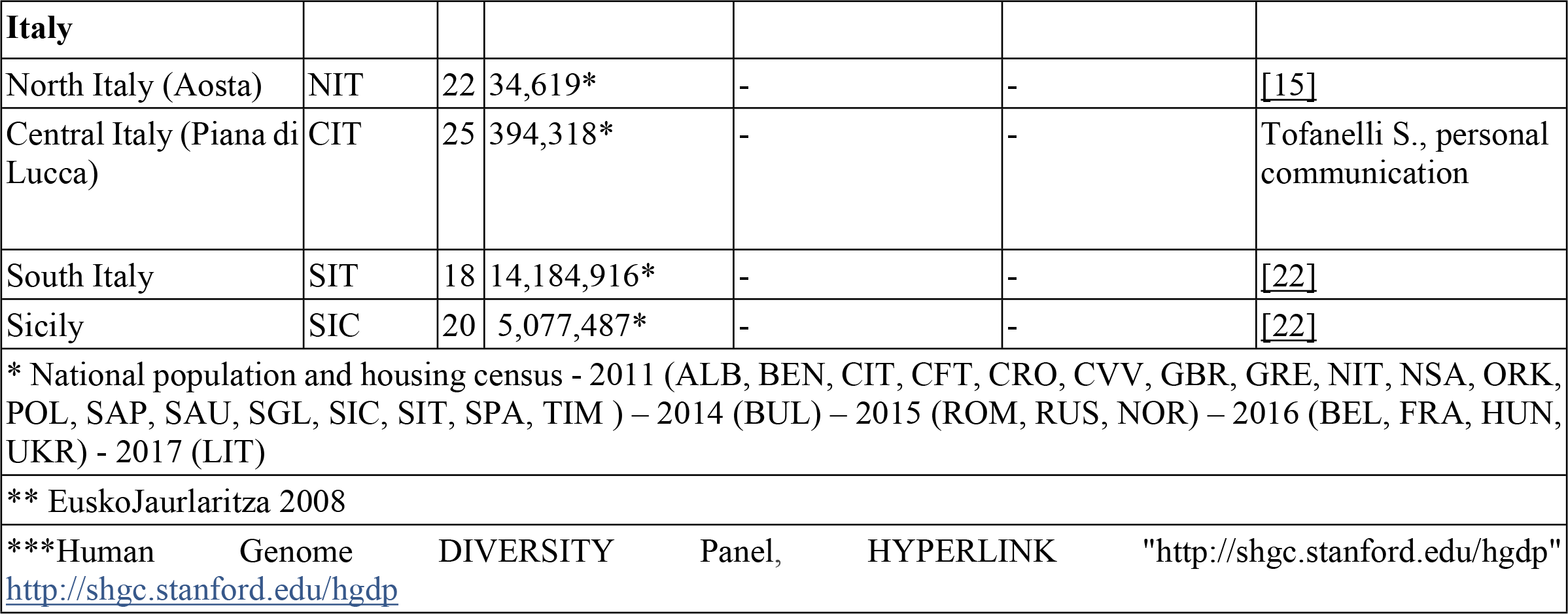
Demographic information about the populations under study.

### Data analyses

The samples genotyped with the GenoChip 2.0 array were merged with literature data and then filtered according to the standard genotype quality control metrics using PLINK (i) SNP genotyping success rate > 90%; (ii) individuals with a genotyping success rate > 92%; (iii) absence of relatedness to the 3rd generation (Identity by Descent, IBD > 0.185). Concerning the latter analysis, when a related pair of individuals was detected, only one sample was randomly chosen and used for the subsequent analysis The PLINK package version 1.9 was used to calculate observed homozygosity (HOM), Identity-by-State (IBS) values, and number (ROH_NSEG) and length (ROH_KB) of Runs of Homozygosity (RoHs). The average HOM per population was estimated using the “--hardy” option. We used the “--distance ibs” option to calculate pairwise intra-population IBS values and calculated the median for each individual’s distribution. The “--homozyg” option was used for RoHs which were identified using the default setting (sliding window of 5 Mb, minimum of 50 SNPs, one heterozygous genotype and five missing calls allowed). In order to ensure that these were true RoHs, we set a minimum-length cut-off of 500 kb and 14 homozygous SNPs [11].

We used SHAPEIT v2.r790 [24] to phase the data, using the 1000 Genomes dataset as a reference panel. We split our dataset by chromosome and phased all individuals simultaneously and used the most likely pairs of haplotypes (using the -output-max option) for each individual for downstream applications. For the phasing and conversion, we used genetic map build 37 downloaded with SHAPEIT. We painted each individual using every other individuals of the same population as a donor [25]. We first inferred the global mutation probability and the switch rate for chromosomes 1, 5, 8, 12, 17 and 22 in 10 iterations of the EM (expectation maximization) algorithm. We fixed the parameters estimated from this analysis (Ne, -n flag, and θ, -M flag) to infer the ChromoPainter coancestry matrix for each chromosome. Using ChromoCombine, we combined the data into a single final coancestry matrix. The haplotype chunks and their total length were estimated using as recipients and donors the individuals of the same population (CHR_P).

The comparison of inter-individual heterogeneity for measures of intra-population variation as well as CHR_P was estimated through the equality of variances (Brown-Forsythe Levene type procedure), after the application of Bonferroni correction (R package lawstat).

Maximum likelihood estimates of individual ancestries were obtained using ADMIXTURE v1.23 under default values. Its algorithm is relatively robust to SNP ascertainment bias [26] since it assigns individual ancestry to a finite number of population clusters, and uses a large multilocus dataset, while the most informative SNPs for ancestry inference are variants with large frequency differences across populations [27]. We applied unsupervised clustering analysis to the whole sample set, exploring the hypothesis of K=2 to 15 clusters. Five independent replicates were run and aligned with CLUMPP. Best K was estimated by the cross-error estimation implemented in ADMIXTURE. We calculated individual heterogeneity (ADX_HET) as the squared difference between each ancestry proportion and its population mean, averaged over all possible ancestries. Population heterogeneity was obtained as the median of individual values.

Admixture dates were inferred using the number of ancestry switches and ancestry proportions following Johnson et al [28]. The whole procedure was as follows: we first jointly phased the 87,818 using the ShapeIt [24] software and the 1000 Genomes data as a reference panel. Phased chromosomes were then used to run the RFMix algorithm [29] with the PopPhased option and default parameters. This modelling approach identifies the ancestry of discrete genomic segments of arbitrary size using a conditional random field parameterized by random forests trained on a reference population panel. Finally, the output of RFmix was employed to calculate both the number of ancestry switches and ancestry proportions for each target individual.

## Results

As a first step, we assessed the genomic heterogeneity occurring among individuals within populations using first four intra-population measures of genomic diversity, based either on single nucleotide (HOM, IBS) or haplotype variation (RoH-KB, RoH-NSEG), for which intra-population variance can be calculated. As a whole, isolated populations showed higher heterogeneity values than the open ones (Fig. 1), with statistically significant differences for two out of four parameters (KB and NSEG; Mann-Whitney test p-value < 0.05). Looking at single populations, the most inbred ones - Sauris, Sappada and Timau - were found to be among the most diverse for all measures along with North Sardinians.

**Fig. 1.**
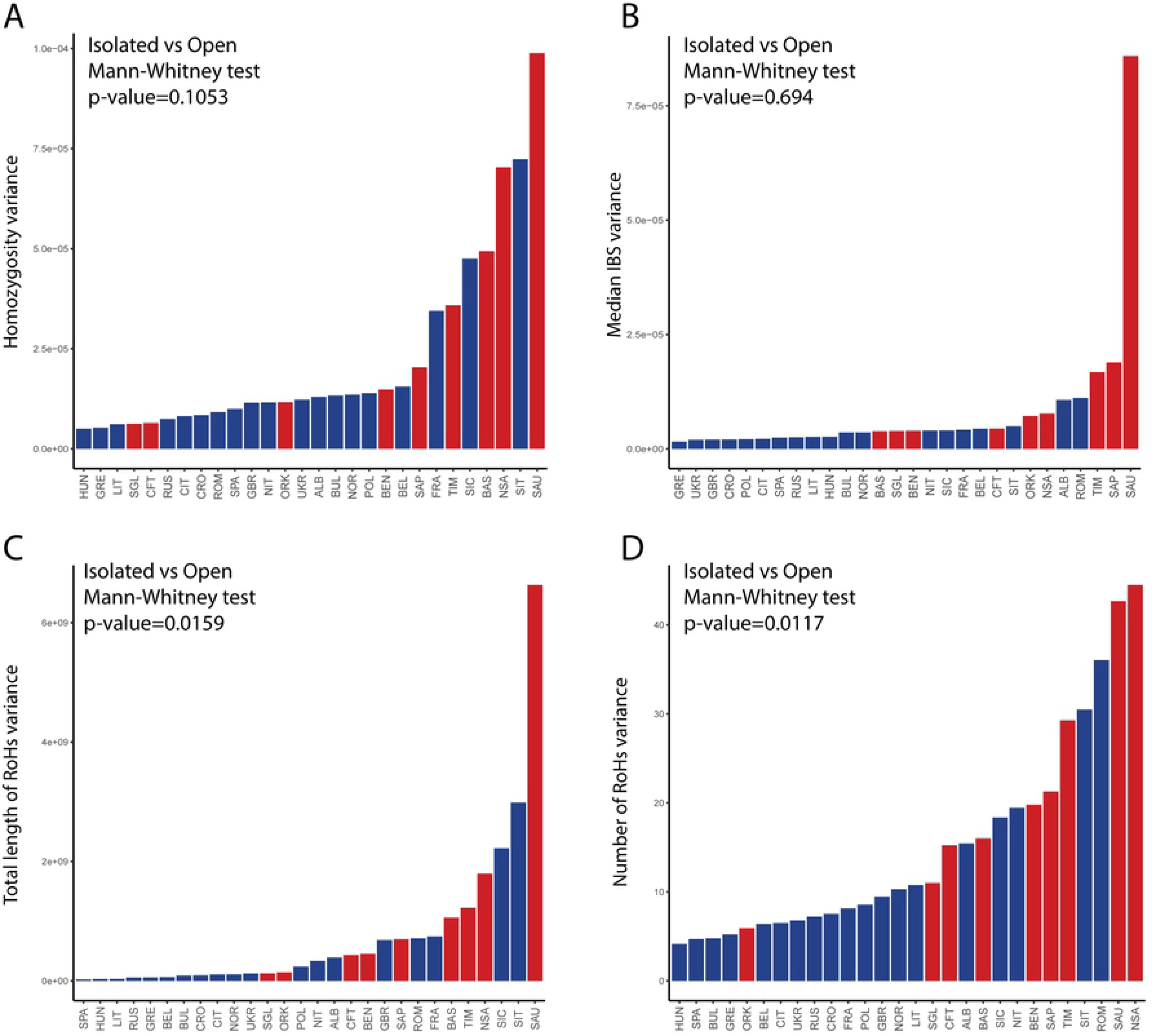
Distribution of inter-individual heterogeneity values across populations and Mann-Whitney U test. Comparison between isolated (red) and open (blue) populations for homozygosity (A), median values of intra-population IBS (B), number of RoHs (C) and total length of RoHs (D).

Then, we compared heterogeneity for ancestry proportions (ADX_HET). Also, in this case, isolates, as a whole, were found to be more heterogeneous than open populations (1.38E-03 vs 6.44E-04), but the difference was statistically insignificant (Mann-U-Whitney p-value > 0.05). The greatest values were again obtained in the three population isolates from the eastern Italian Alps, followed by North Sardinians (Figs 2A and 2B), with a noticeable difference: the heterogeneity was more evenly distributed across individuals of the former populations, as indicated by their ratios between average and median values for the best supported K value (K=4; S1 Tabòe). Interestingly, we detected a highly prevalent village-specific component in 50% of the genomes from Sappada (12 out of 24, at K=4) and in 54% of those from Timau (13 out of 24 at K=5, S1 Fig.). The remaining genomes were clearly more heterogeneous, a likely signature of recent admixture.

**Fig. 2.**
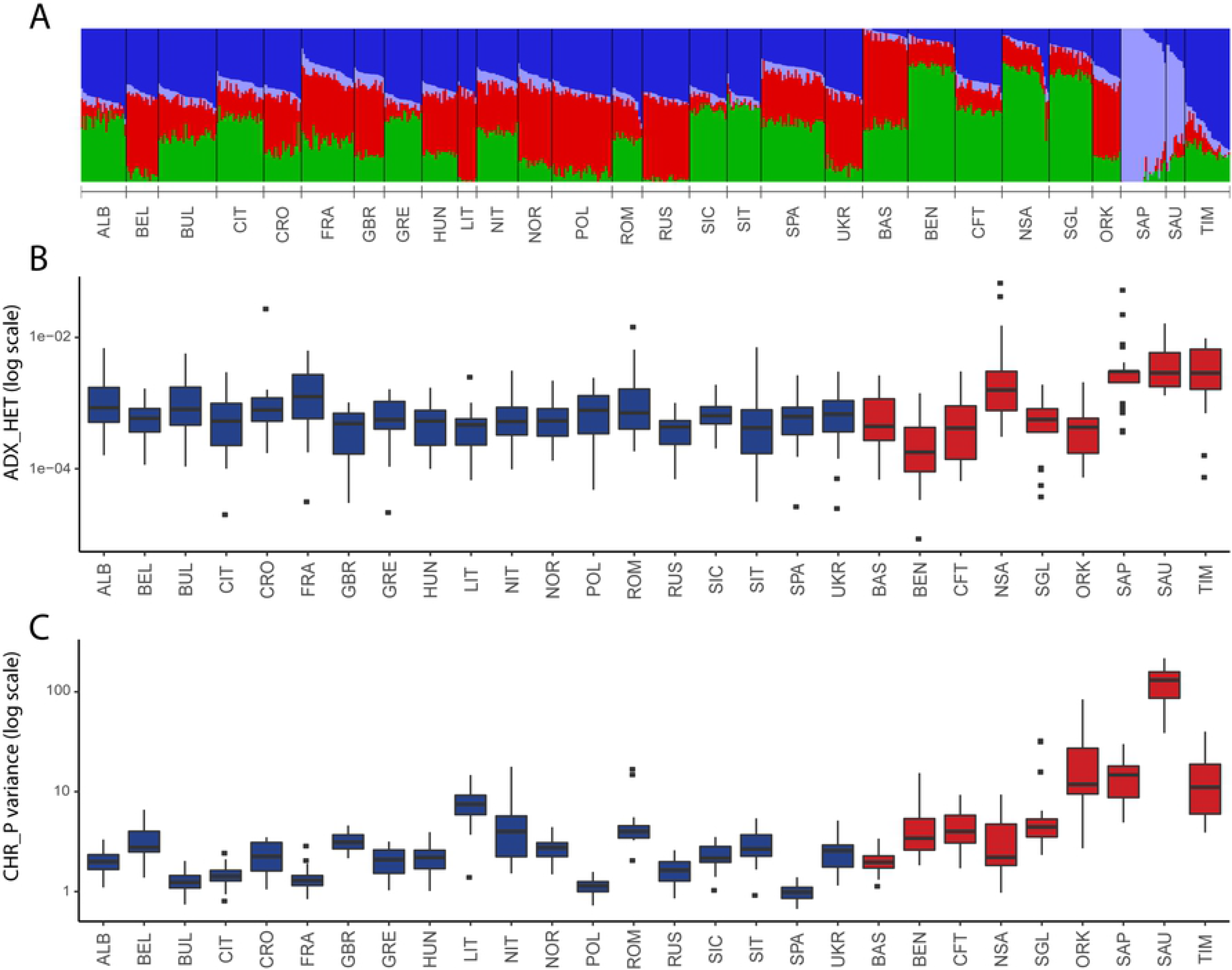
Inter-individual heterogeneity of ancestry components and intra-population haplotype sharing. (A) Maximum likelihood estimates of individual ancestries (K=4) for the 28 populations under study; (B) intra-population distribution of the admixture heterogeneity measure (y axis log scale); (C) Inter-individual heterogeneities of the total length of chunks among individuals in each population (y axis log scale; see materials and methods for more detail).

Finally, we took into account the heterogeneity of the total length of haplotype chunks shared between individuals (CHR_P). The distribution of this parameter reconfirmed the patterns observed for groups (higher values in isolates than open; Mann-Whitney U test based on median variance values, p-value=0.0029) and single populations (higher values in Sauris, Sappada and Timau). As the only peculiarity, a noticeable signal was provided also from the Orkney islanders (Fig. 2C).

In order to understand if the results obtained for the three north eastern Italian isolates might be due to introgression of exogenous genetic components, Sappada and Timau samples were splitted into two sub-groups on the basis of ADMIXTURE ancestry proportions (at K=4 and K=5 for Sappada and Timau, respectively). In the case of Sauris, sub-groups would had been too small to be separately analyzed. Individuals with a highly prevalent village-specific ancestry (threshold 99%; sub-groups SAP_VSA and TIM_VSA) were taken separate from those with more heterogeneous ancestry, who were termed as SAP_HTA and TIM_HTA. Thereafter, we performed the Levene’s tests for equality of variances between all populations (27 comparisons for all combinations population/measure). Only comparisons with a ratio between standard deviations >1 and significant after Bonferroni correction are shown in Fig. 3. The highest number of overall significant comparisons was found for Sauris, which was also the only population with hits in all measures, while the high values of inter-individual heterogeneity for the other north-eastern Italian isolates were not captured by HOM. A relatively high number of significant comparisons still persisted in the HTA groups of both Sappada and Timau, mainly due to KB and CHR_P, respectively. Signatures of inter-individual heterogeneity were recorded also in VSA sub-groups, more evidently in Timau where significant comparisons were observed not only for CHR_P (like in Sappada) but also for KB.

**Fig. 3.**
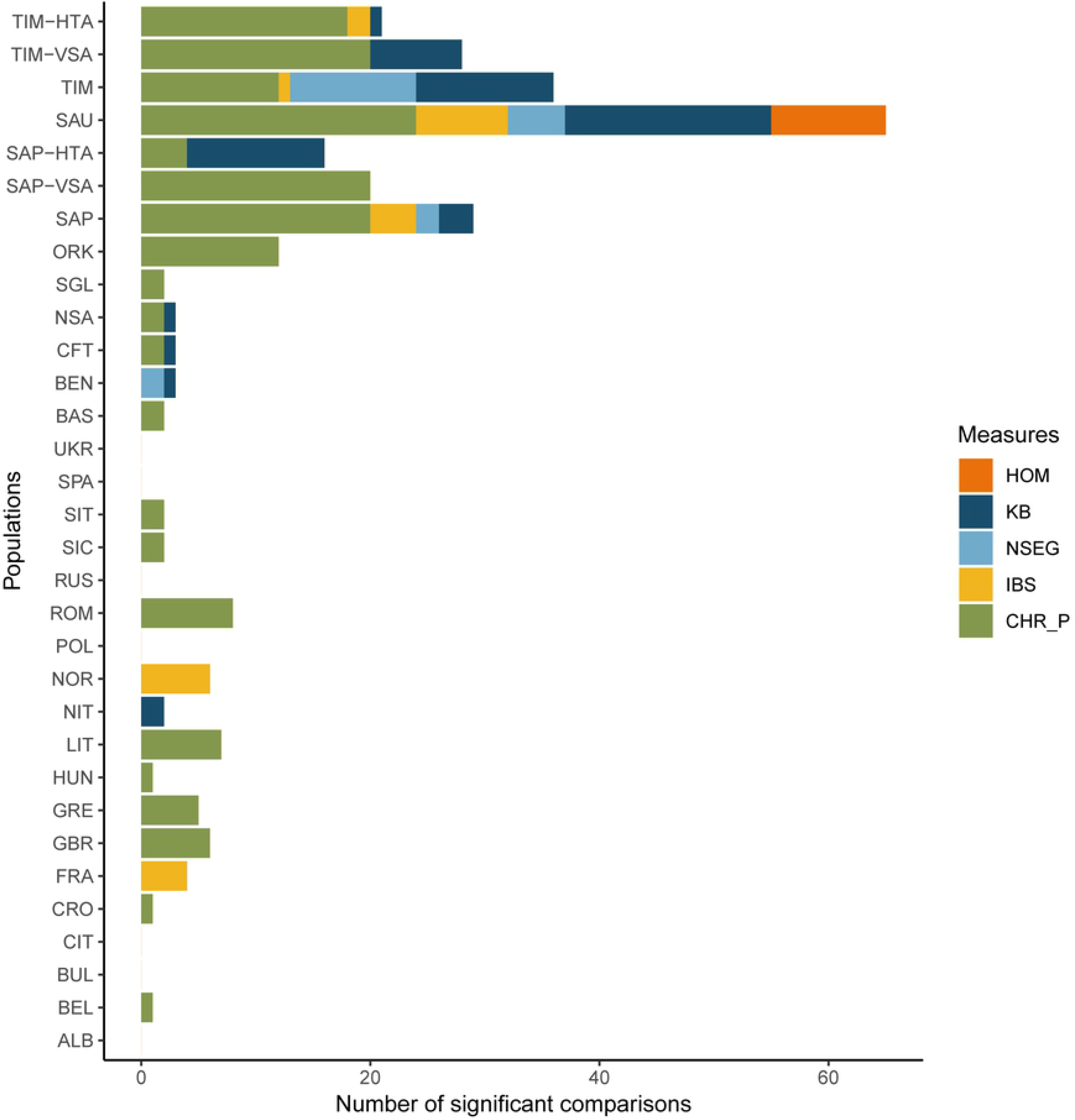
Pairwise comparisons of inter-individual heterogeneity. Number of statistically significant pairwise comparisons with a ratio between standard deviations >1 after Bonferroni correction. For the measures based on pairwise comparisons (IBS and CHR_P), population variance was calculated using the individual median values. Comparisons between Sappada and Timau and their sub-groups (SAP_VSA, SAP_HTA, TIM_VSA and TIM_HTA) were not included.

Given the support received by genetic introgression in generating the observed pattern from the analyses described above, we went to infer the time frames of the admixture which likely occurred between SAP_HTA and TIM_HTA sub-groups and geographically-close Italian speaking populations. We preliminarily tested the reliability of our estimates panel using genomic profiles of African-Americans obtained with a much denser SNP set. To this purpose, we retrieved data from the 1000 genomes project phase 3 and used a simple three population model with 30 randomly chosen individuals from the African-American population (ASW) as targets and an equal number of individuals of European (CEU) and African (YRI) origin as sources. Estimates obtained by using our SNP panel and another including 8,142,382 markers (with MAF<0.05) were close each other and consistent with previous results based on molecular data [30]: the admixture event dated at around six generations ago, with an average value across individuals of 6.9+/−3.7 and 6.2+/−2.8 for the high- and low-density SNP sets, respectively (see S2 Table for individual estimates). Then, we applied the same procedure to the admixed sub-groups (SAP_HTA and TIM_HTA) as targets, while the un-admixed ones (SAP_VSA and TIM_VSA) and the northern Italians (NIT) served as sources. The resulting admixture dates were relatively recent, but consistent with the grandfather rule: from 3.8 to 5.5 generations (average = 4.6) in Sappada and from 3.8 to 4.8 in Timau (average = 4.4) (see S3 and S4 Tables for individual results). As a matter of fact, our sample selection criteria proved effective in avoiding sampling of recently admixed individuals, thereby allowing us to draw a picture of the genomic structure preceding the isolation breakdown, an event occurred in the eastern Alps region between the two world wars [31,32].

## Discussion

Previous GWA studies, which analyzed genetic variation of isolated human populations, focused on measures which summarize single nucleotide and haplotype variation within or among groups [e.g. 11,33,34]. A previous study led by one of us (V.C.) provided evidence of structure within an isolated population (Cardile, southern Italy [35]), but no comparison with other isolates and open populations was carried out. The possible presence of structure within population isolates is worth exploring in depth since it could be a signature of events of recent admixture and/or subdivision; both could potentially disrupt the homogeneity due to the founder effect and persistence of inbreeding over generations.

To gain new insights into the genomic structure of isolated populations, we decided to focus on the distribution of variance (heterogeneity) of intra-population diversity measures across individuals within populations, rather than relying on their average values. In contrast with their common view as groups of genetically homogeneous individuals, we observed that the inter-individual genomic heterogeneity of isolated populations is at least comparable to that of the open ones. It is worth reminding that applying standard measures of intra-population diversity to our dataset produced the expected pattern, with isolates characterized by higher homozygosity, longer and more numerous ROHs and higher IBS values than open populations, although a clear discontinuity of values between the two groups is not noticeable (see [15])

Interestingly, three small and highly inbred isolates (Sappada, Sauris and Timau) were characterized by particularly high heterogeneity values, which largely exceeded those calculated in all other populations. Given that there is no evidence to support the presence of sub-groups with distinct matrimonial behaviours for any of them, this finding could hardly be put down to population subdivision. However, the observed patterns could be explained, at least in part, by relatively recent events of genetic introgression, such as those suggested by our admixture dates based on ancestry switches. In fact, after removing the individuals with higher percentages of mixed ancestries from the Sappada and Timau samplings, their number of statistically significant pairwise comparisons for inter-individual heterogeneity diminished substantially (Fig. 3). We reason that exogenous components might have survived more easily in the three isolates from northeastern Italy than in other populations for two reasons. Firstly, when most, if not all, matrimonial unions occur within small and highly inbred isolates, as is the case for the three populations cited above, carriers of new genetic components may have a greater chance of contributing to the gene pool. In line with this idea, in our global dataset, a high and significant positive correlation was observed between inbreeding rates (S5 Table) and Admixture inter-individual heterogeneity values (Pearson correlation coefficient: 0.768; p-value<0.001). Secondly, the ratio between sample and census size for Sauris, Sappada and Timau (from 1.8% to 4.8%) is greater than in other isolates (from 1.3% to < 0.1%), which increases the probability of sampling individuals bearing genetic components occurring at low or moderate frequencies.

A retrospective look at previous studies shows that other small-sized European isolates with a very high ratio between sample and census size, namely Clauzetto, Erto, Illeggio, Resia and (another sampling from) Sauris, show a similar pattern to what we observed [34]. A high level of heterogeneity among individuals was in fact evidenced by their ancestry proportions and by the results of different types of principal component analyses (basic, spatial and discriminant). The results obtained were explained by Esko et al. [34] as a signature of population sub-structure. Unfortunately, the data this research work was based on were not released by the authors and, therefore, it was not possible to re-analyze and compare them with our results.

Whatever the cause of this high genomic inter-individual heterogeneity we observed in Sappada, Sauris and Timau, we cannot ignore the question: “what do our results imply for the way in which bio-medical studies are carried out in population isolates?”. Although, the most robust evidence was noticed in some young and small-sized population isolates - which are less used in association studies than the older and larger ones [36] - our results are worthy of attention since they highlight a confounding factor which has not been yet adequately taken into account. In fact, to the best of our knowledge, the effect of increased allelic and haplotypic heterogeneity has been investigated only in relation to the issue of undetected population structure in large scale association studies [37], whereas we argue that it may represent a drawback also for genetic investigations of population isolates.

We suggest that genetic clustering algorithms may be used to test for the presence of individuals with different ancestry proportions within isolated populations, similarly to what has been previously done by Esko et al. [29] (see also [38]). Whenever genomes with substantially more heterogeneous ancestry are detected, it would be worth removing them, re-estimating the parameters of gene-disease association and comparing the new results with those obtained using the whole sample. This could help evaluate whether the genomes with mixed ancestry - in which the reduction of the haplotypic and allelic diversity produced by the effects of the founders and inbreeding should be less detectable - may have acted as confounding factors. For each dataset, different ancestry proportions could be tried as thresholds, and the one able to reduce inter-individual heterogeneity without leading to a significant loss of power should be used.

## Conclusions

In this study we have shed light on the occurrence of relatively high levels of inter-individual heterogeneity in populations isolates and proposed a way to monitor their effects on the inferences of association between genes and diseases. This research work challenges the traditional paradigm which considers population isolates as genetically uniform entities, providing evidence of their emerging complexity. We hope that our study can stimulate further investigations based on a wider variety of samples and more powerful genomic tools, through which a better understanding of the fine-grained genomic structure of human population isolates will finally be reached.

## Acknowledgments

We are greatly indebted to all the blood donors. We would also like to thank Marcella Benedetti (Municipality of Sappada), Nino Pacilè and Lucia Protto (Municipality of Sauris), Vito Massalongo (Giazza), Ottaviano Matiz and Velia Plozner (Timau) for their valuable assistance in the sample collection and for their warm hospitality.

## Supporting information

**S1 Table. Ratio between mean and median inter-individual heterogeneity**. Analysis based on the Admixture components proportion recorded at K=4.

**S2 Table. Date estimates based on ancestry switches inferred with the high- and low-density SNP sets for the 1000 Genomes African Americans**.

**S3 Table. Ancestry proportions, number of ancestry switches and date estimates for the Sappada admixed subgroup**.

**S4 Table. Ancestry proportions, number of ancestry switches and date estimates for the Timau admixed subgroup**.

**S5 Table. Inbreeding coefficient values**. Calculated as the proportion of the autosomal genome in runs of homozygosity, excluding the centromeres.

**S1 Fig. Maximum likelihood estimates of individual ancestries**. Plots from K=2 to K=10 for the 28 populations under study.

## References

1. Ward RH, Neel JV. Gene frequencies and microdifferentiation among the Makiritare Indians. IV. A comparison of a genetic network with ethnohistory and migration matrices; a new index of genetic isolation. Am J Hum Genet. 1970;22: 538–561. Available: https://www.ncbi.nlm.nih.gov/pubmed/5516237

2. Arcos-Burgos M, Muenke M. Genetics of population isolates. Clin Genet. 2002;61: 233–247. doi:10.1034/j.1399-0004.2002.610401.x

3. Colonna V, Nutile T, Astore M, Guardiola O, Antoniol G, Ciullo M, et al. Campora: a young genetic isolate in South Italy. Hum Hered. 2007;64: 123–135. doi:10.1159/000101964

4. Charlesworth B. Fundamental concepts in genetics: effective population size and patterns of molecular evolution and variation. Nat Rev Genet. 2009;10: 195–205. doi:10.1038/nrg2526

5. Palin K, Campbell H, Wright AF, Wilson JF, Durbin R. Identity-by-descent-based phasing and imputation in founder populations using graphical models. Genet Epidemiol. 2011;35: 853–860. doi:10.1002/gepi.20635

6. de la Chapelle A, Wright FA. Linkage disequilibrium mapping in isolated populations: The example of Finland revisited. Proceedings of the National Academy of Sciences. 1998;95: 12416–12423. doi:10.1073/pnas.95.21.12416

7. Jorde LB, Watkins WS, Kere J, Nyman D, Eriksson AW. Gene mapping in isolated populations: new roles for old friends? Hum Hered. 2000;50: 57–65. doi:10.1159/000022891

8. Varilo T, Laan M, Hovatta I, Wiebe V, Terwilliger JD, Peltonen L. Linkage disequilibrium in isolated populations: Finland and a young sub-population of Kuusamo. Eur J Hum Genet. 2000;8: 604–612. doi:10.1038/sj.ejhg.5200482

9. Service S, DeYoung J, Karayiorgou M, Roos JL, Pretorious H, Bedoya G, et al. Magnitude and distribution of linkage disequilibrium in population isolates and implications for genome-wide association studies. Nat Genet. 2006;38: 556–560. doi:10.1038/ng1770

10. Kristiansson K, Naukkarinen J, Peltonen L. Isolated populations and complex disease gene identification. Genome Biol. 2008;9: 109. doi:10.1186/gb-2008-9-8-109

11. Colonna V, Pistis G, Bomba L, Mona S, Matullo G, Boano R, et al. Small effective population size and genetic homogeneity in the Val Borbera isolate. Eur J Hum Genet. 2013;21: 89–94. doi:10.1038/ejhg.2012.113

12. Hatzikotoulas K, Gilly A, Zeggini E. Using population isolates in genetic association studies. Brief Funct Genomics. 2014;13: 371–377. doi:10.1093/bfgp/elu022

13. Panoutsopoulou K, Hatzikotoulas K, Xifara DK, Colonna V, Farmaki A-E, Ritchie GRS, et al. Genetic characterization of Greek population isolates reveals strong genetic drift at missense and trait-associated variants. Nat Commun. 2014;5: 5345. doi:10.1038/ncomms6345

14. Xue Y, Mezzavilla M, Haber M, McCarthy S, Chen Y, Narasimhan V, et al. Enrichment of low-frequency functional variants revealed by whole-genome sequencing of multiple isolated European populations. Nat Commun. 2017;8: 15927. doi:10.1038/ncomms15927

15. Anagnostou P, Dominici V, Battaggia C, Pagani L, Vilar M, Wells RS, et al. Overcoming the dichotomy between open and isolated populations using genomic data from a large European dataset. Sci Rep. 2017;7: 41614. doi:10.1038/srep41614

16. Elhaik E, Greenspan E, Staats S, Krahn T, Tyler-Smith C, Xue Y, et al. The GenoChip: a new tool for genetic anthropology. Genome Biol Evol. 2013;5: 1021–1031. doi:10.1093/gbe/evt066

17. Capocasa M, Anagnostou P, Bachis V, Battaggia C, Bertoncini S, Biondi G, et al. Linguistic, geographic and genetic isolation: a collaborative study of Italian populations. J Anthropol Sci. 2014;92: 201–231. doi:10.4436/JASS.92001

18. Anagnostou P, Capocasa M, Dominici V, Montinaro F, Coia V, Destro-Bisol G. Evaluating mtDNA patterns of genetic isolation using a re-sampling procedure: A case study on Italian populations. Ann Hum Biol. 2017;44: 140–148. doi:10.1080/03014460.2016.1181784

19. Human Genome Diversity Project (HGDP). Encyclopedia of Genetics, Genomics, Proteomics and Informatics. 2008. pp. 923–923. doi:10.1007/978-1-4020-6754-9_7923

20. Sarno S, Boattini A, Pagani L, Sazzini M, De Fanti S, Quagliariello A, et al. Ancient and recent admixture layers in Sicily and Southern Italy trace multiple migration routes along the Mediterranean. Sci Rep. 2017;7: 1984. doi:10.1038/s41598-017-01802-4

21. Behar DM, Yunusbayev B, Metspalu M, Metspalu E, Rosset S, Parik J, et al. The genome-wide structure of the Jewish people. Nature. 2010;466: 238–242. doi:10.1038/nature09103

22. Hellenthal G, Busby GBJ, Band G, Wilson JF, Capelli C, Falush D, et al. A genetic atlas of human admixture history. Science. 2014;343: 747–751. doi:10.1126/science.1243518

23. Yunusbayev B, Metspalu M, Järve M, Kutuev I, Rootsi S, Metspalu E, et al. The Caucasus as an asymmetric semipermeable barrier to ancient human migrations. Mol Biol Evol. 2012;29: 359–365. doi:10.1093/molbev/msr221

24. O’Connell J, Gurdasani D, Delaneau O, Pirastu N, Ulivi S, Cocca M, et al. A general approach for haplotype phasing across the full spectrum of relatedness. PLoS Genet. 2014;10: e1004234. doi:10.1371/journal.pgen.1004234

25. Lawson DJ, Hellenthal G, Myers S, Falush D. Inference of population structure using dense haplotype data. PLoS Genet. 2012;8: e1002453. doi:10.1371/journal.pgen.1002453

26. Haasl RJ, Payseur BA. Multi-locus inference of population structure: a comparison between single nucleotide polymorphisms and microsatellites. Heredity. 2011;106: 158–171. doi:10.1038/hdy.2010.21

27. Rosenberg NA, Li LM, Ward R, Pritchard JK. Informativeness of genetic markers for inference of ancestry. Am J Hum Genet. 2003;73: 1402–1422. doi:10.1086/380416

28. Johnson NA, Coram MA, Shriver MD, Romieu I, Barsh GS, London SJ, et al. Ancestral Components of Admixed Genomes in a Mexican Cohort. PLoS Genet. 2011;7: e1002410. doi:10.1371/journal.pgen.1002410

29. Maples BK, Gravel S, Kenny EE, Bustamante CD. RFMix: a discriminative modeling approach for rapid and robust local-ancestry inference. Am J Hum Genet. 2013;93: 278–288. doi:10.1016/j.ajhg.2013.06.020

30. Moorjani P, Patterson N, Hirschhorn JN, Keinan A, Hao L, Atzmon G, et al. The history of African gene flow into Southern Europeans, Levantines, and Jews. PLoS Genet. 2011;7: e1001373. doi:10.1371/journal.pgen.1001373

31. Vogel F. Break-up of isolates. In: Roberts DF, Fujiki N, Torizuka K, Roberts DF, Fujiki N, Torizuka K, editors. Isolation, Migration and Health. Cambridge: Cambridge University Press; 1992. pp. 41–54. doi:10.1017/CBO9780511983634.006

32. Viazzo PP. Transizioni alla modernità in area alpina. Dicotomie, paradossi, questioni aperte. Histoire des Alpes – Storia delle Alpi – Geschichte der Alpen 2007;12: 13–28.

33. Karafet TM, Bulayeva KB, Bulayev OA, Gurgenova F, Omarova J, Yepiskoposyan L, et al. Extensive genome-wide autozygosity in the population isolates of Daghestan. Eur J Hum Genet. 2015;23: 1405–1412. doi:10.1038/ejhg.2014.299

34. Esko T, Mezzavilla M, Nelis M, Borel C, Debniak T, Jakkula E, et al. Genetic characterization of northeastern Italian population isolates in the context of broader European genetic diversity. Eur J Hum Genet. 2013;21: 659–665. doi:10.1038/ejhg.2012.229

35. Colonna V, Nutile T, Ferrucci RR, Fardella G, Aversano M, Barbujani G, et al. Comparing population structure as inferred from genealogical versus genetic information. Eur J Hum Genet. 2009;17: 1635–1641. doi:10.1038/ejhg.2009.97

36. Heutink P, Oostra BA. Gene finding in genetically isolated populations. Hum Mol Genet. 2002;11: 2507–2515. Available: https://www.ncbi.nlm.nih.gov/pubmed/12351587

37. Marchini J, Cardon LR, Phillips MS, Donnelly P. The effects of human population structure on large genetic association studies. Nat Genet. 2004;36: 512–517. doi:10.1038/ng1337

38. Rosenberg NA, Pritchard JK, Weber JL, Cann HM, Kidd KK, Zhivotovsky LA, et al. Genetic structure of human populations. Science. 2002;298: 2381–2385. doi:10.1126/science.1078311

